# Retrotransposon-mediated duplication of *SSU1* in high SO_2_ tolerant *Brettanomyces bruxellensis* winery isolates

**DOI:** 10.1101/2025.10.09.681542

**Authors:** Cristobal A. Onetto, Joseph Rossi, Antonio G. Cordente, Steven Van Den Heuvel, Anthony R. Borneman

## Abstract

*Brettanomyces bruxellensis* is an industrially relevant yeast and major spoilage organism in wine, where tolerance to sulphur dioxide (SO_2_), the primary preservative used for its control, varies between strains. To investigate the genetic basis of SO_2_ tolerance in winery populations, 26 isolates from Australian wineries were phenotypically characterised and sequenced using long-read technology. Isolates showed a wide range of SO_2_ tolerance that correlated with phylogenetic clade and ploidy. Haplotype phasing of *SSU1*, a sulphite efflux pump linked to SO_2_ tolerance, identified nine distinct haplotypes, including the previously described high-tolerance H1 allele. Highly tolerant strains carried duplications of H1, frequently associated with retrotransposon insertions and chromosomal rearrangements at the *SSU1* locus. Comparative analyses with laboratory-evolved strains confirmed that retrotransposons facilitated the acquisition of additional *SSU1* copies. These findings suggest that transposon-mediated structural variation drives adaptive increases in SO_2_ tolerance in *B. bruxellensis* populations.

## 1. Introduction

*Brettanomyces bruxellensis* is a commercially important yeast associated with the production of wine, beer, and cider [1]. Its physiological traits, including tolerance to low pH and high ethanol, a broad carbon-utilisation profile, and, in some strains, the ability to assimilate nitrate, confer competitive advantages over other yeasts commonly found in fermented beverages [1-3]. In wine, *B. bruxellensis* is generally regarded as a spoilage organism because it can convert hydroxycinnamic acids to 4-ethylphenol and 4-ethylguaiacol, compounds associated with ‘horsey’, ‘barnyard’, ‘smoky’, and ‘band-aid’ aromas that also alter the wine’s flavour profile [2, 4].

To inhibit the growth of *B. bruxellensis*, winemakers commonly add sulfur dioxide (SO_2_) at the end of fermentation. Beyond its antioxidant activity, SO_2_ suppresses yeast growth by interfering with key intracellular processes [5]. *B. bruxellensis* shows considerable strain variability in SO_2_ tolerance [6, 7], which has been linked to specific polyploid genotypes frequently isolated from wine [8]. Consequently, the routine use of SO_2_ in wineries is thought to lead to selection for SO_2_-tolerant genotypes [6, 8, 9].

A major mechanism of sulphite tolerance in yeasts is active efflux mediated by a plasma-membrane pump encoded by the gene *SSU1* [10]. In *Saccharomyces cerevisiae*, highly sulphite-tolerant strains often carry chromosomal rearrangements that place *SSU1* under a stronger promoter (a translocation generating an SSU1-ECM34 promoter fusion), resulting in elevated *SSU1* expression and increased tolerance to SO_2_ [11-13]. *B. bruxellensis* also harbours an *SSU1* ortholog [14]. Deleting *SSU1* in a haploid background reduces sulphite tolerance, even at very low concentrations [15]. Additionally, a previous study linked particular *B. bruxellensis SSU1* haplotypes with higher tolerance to SO_2_ [14]. Under sustained SO_2_ exposure in laboratory evolution, genomic changes arise that increase the copy number of toleran*t SSU1* haplotypes [9]. Together, these findings indicate that *SSU1* is a major contributor to sulphite tolerance in *B. bruxellensis*, although recent work suggests additional loci and mechanisms may be also linked to sulphite tolerance in this species [7].

Although laboratory evolution shows that polyploid *B. bruxellensis* can rapidly acquire increased SO_2_ tolerance through genomic variation and *SSU1* duplication [9], it remains unclear whether SO_2_-tolerance is occurring in the field through similar genetic adaptations. To address this, *B. bruxellensis* strains were isolated from wineries across Australia and then subjected to both phenotypic characterisation of SO_2_ tolerance and detailed genomic comparison to investigate the genetic basis of SO_2_ tolerance across industry isolates.

## 2. Material and Methods

### 2.1 Brettanomyces bruxellensis isolates

Wineries in South Australia provided samples suspected to have *Brettanomyces* contamination. Selective isolation of *B. bruxellensis* from the samples was achieved by using Wallerstein Laboratories (WL) nutrient agar (Millipore) supplemented with cycloheximide (100 mg/L), chloramphenicol (50 mg/L) and biphenyl (500 mg/L) (WL + CCB). Each sample was processed in one of the following ways: (a) 10-50 mL of wine was filtered through a 0.45 µM filter and the membrane was placed on the WL + CCB media, or (b) 1 mL of wine was concentrated down to a final volume of 100 µL, and the entire 100 µL was aseptically spread-plated on the WL + CCB media. Plates were then incubated for at least 7 days at 28 °C in the dark to allow for *B. bruxellensis* colonies to grow. Confirmation of isolates as *B. bruxellensis* was confirmed through ITS by comparing the product length with a *B. bruxellensis* control.

### 2.2 Sulphite tolerance phenotyping

Sulphite tolerance screening was performed as previously described in Bartel, Roach (9) with minor modifications. Briefly, isolates were inoculated in quadruplicate into 96 deep-well plates containing 800 µL of sterile minimal medium (MM) containing 6.7 g/L of yeast nitrogen base with amino acids and 5 g/L of glucose at pH 3.5. Strains were incubated at 28 °C for 3 days until they reached saturation and then 10 µL was sub-cultured into 190 µL of MM supplemented with a freshly prepared, filter sterilised, potassium metabisulphite solution (PMS) (20 g/L) to a final molecular SO_2_ of 0, 0.098, 0.223, 0.335, 0.446, 0.558, 0.669 and 0.781 mg/L. Plates were incubated at 28 °C for 6 days and then OD was measured at 600 nm. Growth was classified as positive if their relative growth to the control condition (OD600 no SO_2_/added SO_2_) was higher than 0.2. Strains were classified into low (growth at < 0.223 mg/L molecular SO_2_), medium (growth between 0.223-0.446 mg/L molecular SO_2_) and high (growth at ≥ 0.558 mg/L molecular SO_2_) SO_2_ tolerant.

### 2.3 DNA extraction and genome sequencing

For long-read sequencing, DNA was extracted using a custom protocol. A 2 mL overnight culture was resuspended in 800 µL of zymolyase solution (0.9 M sorbitol, 0.1 M EDTA, 16.2 µL zymolyase [10 mg/mL], 10 mM DTT) and incubated under agitation at 37 °C for 1 h. Cells were pelleted and resuspended in 400 µL of lysis buffer (0.2 M LiAc, 1% SDS, 100 mM Tris-HCl pH 7.5, 50 mM EDTA) and incubated at 75 °C for 30 min with agitation at 800 rpm. Samples were combined with an equal volume of 0.1 mm and 0.5 mm silica/zirconia beads and subjected to bead beating (8000 rpm, 6 × 30 s cycles with 30 s pauses). Next, 100 µL of 5 M KAc was added to the lysate, mixed, and incubated on ice for 10 min. Samples were centrifuged, and the supernatant was mixed with 1 mL of ethanol and centrifuged at maximum speed for 4 min. The resulting DNA pellet was washed with 70 % ethanol, resuspended in 100 µL of TE buffer (Invitrogen, Cat. 12090-015) containing 1 µL of RNase A (100 µg/mL), and incubated at 37 °C for 30 min.

Sequencing libraries were prepared using the SQK-NBD114-24 kit and sequenced in FLO-PRO114M flow cell (Oxford Nanopore Technologies, Oxford, UK). Pod5 files were base-called using Dorado (https://github.com/nanoporetech/dorado) with the Sup model v4.3.0.

### 2.4 Genome assembly, annotation and SSU1 haplotype phasing

Ploidy estimations were performed using Smudgeplot v. 0.4.0 [16] and manual allele depth observations. Long-read haplotype resolved assemblies were performed using hifiasm v. 0.25 [17] and polished using Medaka (https://github.com/nanoporetech/medaka). Gene prediction was performed following the funannotate pipeline [18].

For *SSU1* phasing, an initial manual assessment of copy number variation (CNV) and amino acid sequences of annotated *SSU1* genes was conducted using the phased genome assembly of each strain. This was cross validated by inspecting read coverage and allele depth at the *SSU1* locus using the integrative genomics viewer (IGV). Local spikes in coverage were observed in strains with duplications of annotated and phased *SSU1* genes. To estimate the copy number of each allele, the major allele frequency at each site was calculated. Additionally, local phasing and reassembly at the *SSU1* locus were confirmed using a combination of FreeBayes v. 1.3.10 [19], with ploidy informed by the manually estimated copy number, and WhatsHap v. 2.8 [20] with the polyphase function. Transposable element (TE) annotation was performed using EDTA v. 2.2.2 [21] and TE insertions identified using the GraffiTE pipeline [22].

## 3. Results and Discussion

Previous investigations have demonstrated substantial variability in the SO_2_ tolerance of wine isolates of *B. bruxellensis*, which was highly correlated with specific phylogenetic clades [6-8]. It has also been demonstrated that this species can adapt to continuous SO_2_ exposure in the laboratory, with strains increasing tolerance through genomic variation, including duplication of the *SSU1* transporter, which forms part of the SO_2_ tolerance mechanism in this species [9].

To attempt extending this link between SO_2_ tolerance and genomic variation to field populations, 26 isolates obtained from wineries across Australia were subjected to detailed SO_2_ phenotyping and whole-genome comparison using long-read sequencing. Phenotyping was carried out by assessing growth in liquid medium across a gradient of increasing SO_2_ concentrations. A large (< 0.223-0.669 mg/L molecular SO_2_) degree of variation in SO_2_ tolerance was observed (Figure 1a), which is consistent with previous global surveys of *B. bruxellensis* [2, 8]. Using this data, three groups were defined (Figure 1a): low tolerance (no growth in 0.223 mg/L SO_2_), intermediate tolerance (growth between 0.223-0.446 mg/L SO_2_) and high tolerance (growth at ≥ 0.558 SO_2_).

**Figure 1.**
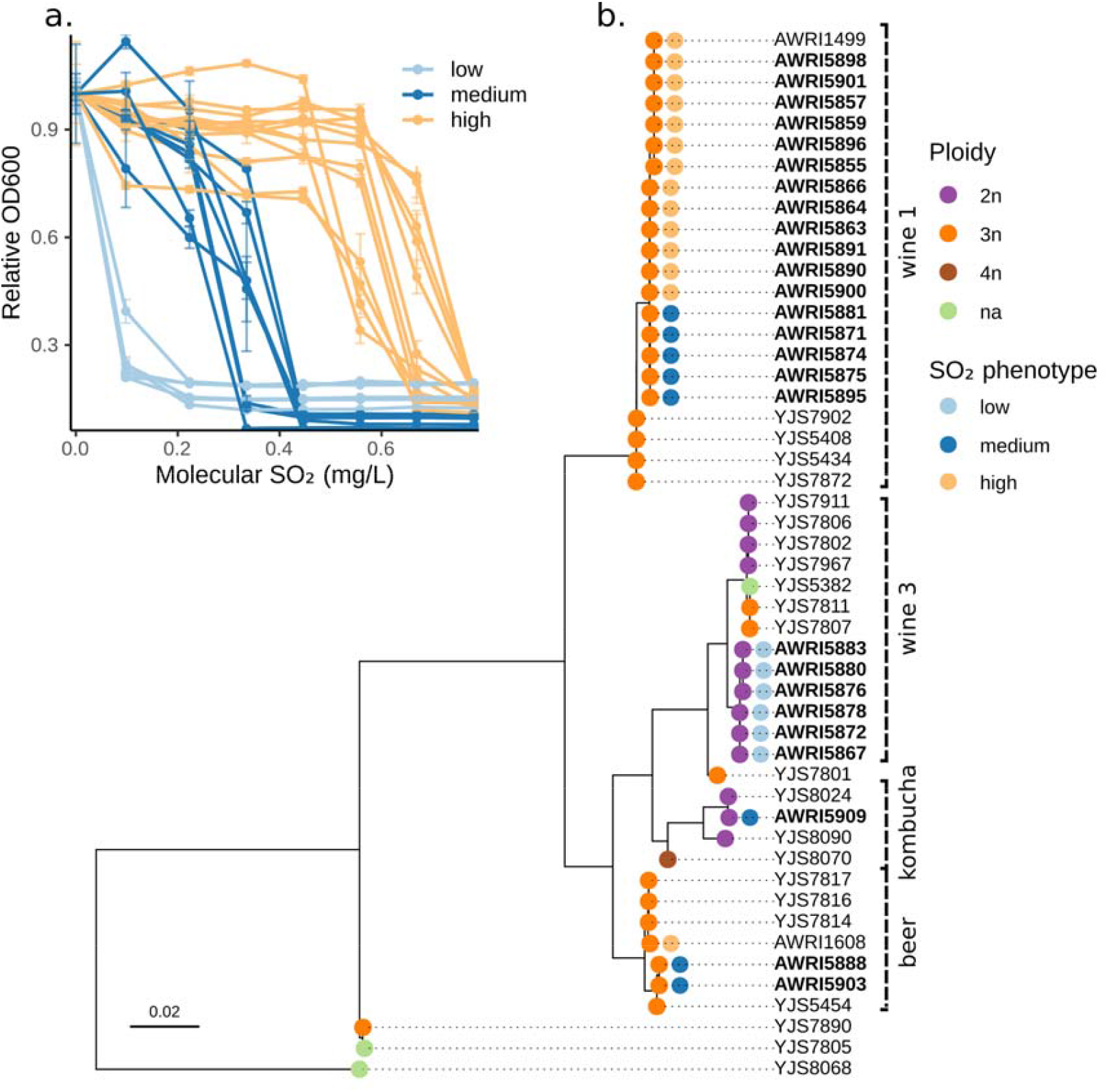
Phenotypic characterisation and phylogeny of 26 *Brettanomyces bruxellensis* isolates. **a.** Sulphite tolerance of *B. bruxellensis* isolates obtained from wineries across Australia. Isolates were grown in quadruplicate in increasing SO_2_ concentrations, and their growth was assessed after 6 days of incubation. Strains were classified into low (growth at < 0.223 mg/L molecular SO_2_), medium (growth between 0.223-0.446 mg/L molecular SO_2_) and high (growth at ≥ 0.558 mg/L molecular SO_2_) sulphite tolerant based on their relative growth to the control condition (OD600 no SO_2_/added SO_2_). **b**. Maximum-likelihood phylogenetic tree constructed from single nucleotide polymorphisms (SNPs) identified using the *B. bruxellensis* reference genome of strain UCD 2041. Isolates presented in this study are highlighted in **bold** and their estimated ploidy and SO_2_ tolerance classification are presented. Selected isolated previously published [23] were also included in the tree reconstruction.

To determine whether these phenotypic categories were associated with specific genotypes, long-read whole-genome sequencing was performed on all 26 isolates. Reads were mapped to the UCD 2041 reference strain genome alongside previously published sequencing data from isolates representing the major *B. bruxellensis* phylogenetic clades [23], with single nucleotide polymorphisms (SNPs) identified for use in phylogenetic reconstruction. The isolates were subsequently assigned to four phylogenetic clades (Figure 1b), including previously defined wine 1 AWRI1499-like (recently renamed A2), wine 3 (recently renamed D1), kombucha (recently renamed D2) and beer (recently renamed A1) clades [7, 23]. As observed in previous studies of Australian *B. bruxellensis* wine populations, most strains were shown to be part of the AWRI1499 allopolyploid wine clade (Figure 1b). All *B. bruxellensis* isolates within this clade were estimated to be triploid (Table S1) and included isolates with medium and high SO_2_ tolerance phenotypes (Figure 1b). Strains within this clade are consistently reported as high SO_2_ tolerant [6, 8], although variability within the clade is also observed [7]. This suggests that genetic variation within this clade is likely responsible for adaptation to higher levels of SO_2_. In contrast, all isolates classified as having low SO_2_ tolerance fell within a separate clade (wine 3) and were identified as diploids (Figure 1b).

### 3.1 Duplication of the *SSU1* SO□-tolerant haplotype in SO□-tolerant *B. bruxellensis* wine industry isolates

The SSU1 sulphite efflux pump has been shown to play a direct role in the tolerance of *B. bruxellensis* to SO_2_ [14, 24]. To investigate the genetic variation underlying differences in SO_2_ tolerance, Varela et al. (2022) examined *SSU1* haplotypes derived from strain AWRI1499, demonstrating that different haplotypes conferred varying levels of tolerance when expressed in *Saccharomyces cerevisiae*. One haplotype (herein referred to as H1), which was distinguished by two unique amino acid residues, was shown to confer the highest SO_2_ tolerance. This same H1 haplotype was duplicated in AWRI1499 strains subjected to continuous SO_2_ exposure during directed evolution, which resulted in increased tolerance after 50 and 100 generations [9]. These laboratory findings suggested a potential mechanism of adaptation in winery isolates. However, it remained unclear whether SO_2_-tolerant isolates sourced directly from winemaking environments also display *SSU1* duplications, and if so, whether these are associated with specific haplotypes. To address these questions, long-read sequencing data were analysed to phase *SSU1* haplotypes and assess CNV across the 26 isolates (Figure 2).

**Figure 2.**
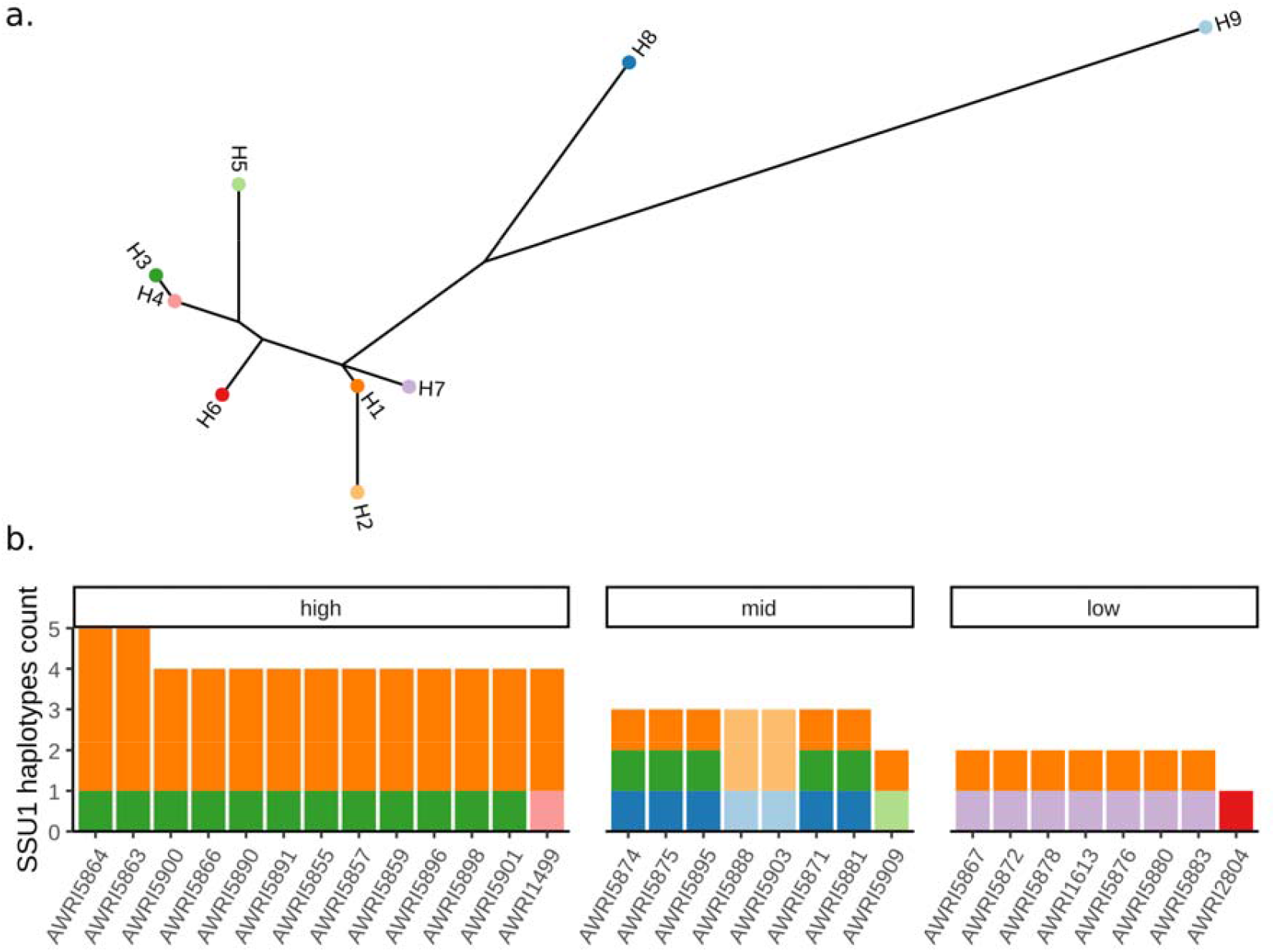
*SSU1* haplotypes in *Brettanomyces bruxellensis*. **a.** Maximum-likelihood phylogenetic tree constructed using the protein sequences of phased *SSU1* haplotypes identified in the long-read genomes of 29 *B. bruxellensis* isolates. H1 represents the *SSU1* haplotype previously shown to confer high SO_2_ tolerance [14] **b**. *SSU1* haplotype count for each of the isolates sequenced using long-reads faceted based on their SO_2_ phenotypes.

Nine distinct protein haplotypes, including previously reported haplotypes H1 and H7 of diploid strain AWRI1613, were detected amongst the *B. bruxellensis* population (Figure 2A). The AWRI1499 H1 haplotype was detected across all three SO_2_ phenotype categories and in both triploid and diploid strains (Figure 2B). Only one additional haplotype (H2), which was identified in two triploid strains, also contained both the D143 and R208 amino acid residues previously implicated in higher SO_2_ tolerance in the AWRI1499 H1 haplotype (Figure S1) [14]. However, H2 differed from H1 by three additional amino acid substitutions (Figure S1). All low SO_2_ tolerant diploid strains contained the SSU1 haplotypes previously reported in the diploid strain AWRI1613 (Figure 2b) [14]

Copy number analysis revealed all high SO_2_-tolerant triploid strains to have at least one duplication of *SSU1*, with two strains possessing five copies each (Figure 2B). In every case, the duplications observed in the high SO_2_-tolerant strains were associated with the H1 haplotype, strongly suggesting that this haplotype is under positive selection. In contrast to previous reports [14], read depth analysis at the *SSU1* locus in AWRI1499 confirmed the presence of four total copies of *SSU1*, three of which corresponded to the H1 haplotype (Figure 2B). In addition, the detection of three distinct *SSU1* haplotypes in medium-tolerant triploid strains belonging to the AWRI1499-like clade suggests that ongoing genetic adaptation within this lineage contributes to increased SO_2_ tolerance. Further work will be required to determine whether the observed increase in tolerance is driven specifically by duplication of the H1 haplotype and/or by *SSU1* gene dosage effects.

### 3.2 Retrotransposon-associated structural variation in the *SSU1* locus

*SSU1* phasing and CNV analyses indicated that SO_2_-tolerant winery isolates carried duplications of a specific *SSU1* haplotype. This phenomenon was also observed in strain AWRI1499 following continuous SO_2_ exposure (Bartel et al. 2021), suggesting that the *SSU1* locus may represent a hotspot for genomic variation. In eukaryotes, repetitive elements are a recognised driver of genomic variation, with repeat-rich regions such as centromeres, sub-telomeres, and transposable element (TE) clusters showing a higher propensity for structural change [25, 26].

To investigate whether repetitive elements within the *SSU1* locus might underlie the observed CNVs, long-read phasing and assembly of the *SSU1* locus was generated for strains of differing ploidy that represented all three SO_2_ tolerance groups (Figure 3). Synteny analyses between haplotypes revealed structural variation (SV) in the *SSU1* locus across strains (Figure 3). Retrotransposon insertions were detected at two positions within the locus, an LTR-Gypsy element located directly downstream of *SSU1* in the triploid strains AWRI5874 and AWRI5891 and an LTR-Copia element upstream of *BEM2* in both diploid and triploid strains (Figure 3). Synteny breakpoints adjacent to the LTR-Copia insertion were observed in triploid strains (Figure 3). Variation in the presence of TE insertions within the *SSU1* locus was evident between clades. All strains belonging to the AWRI1499 clade contained TE insertions, whereas none of the Beer or Kombucha clade strains contained insertions. In the Wine 3 clade, three strains carried TE insertions while three did not (Table S1).

**Figure 3.**
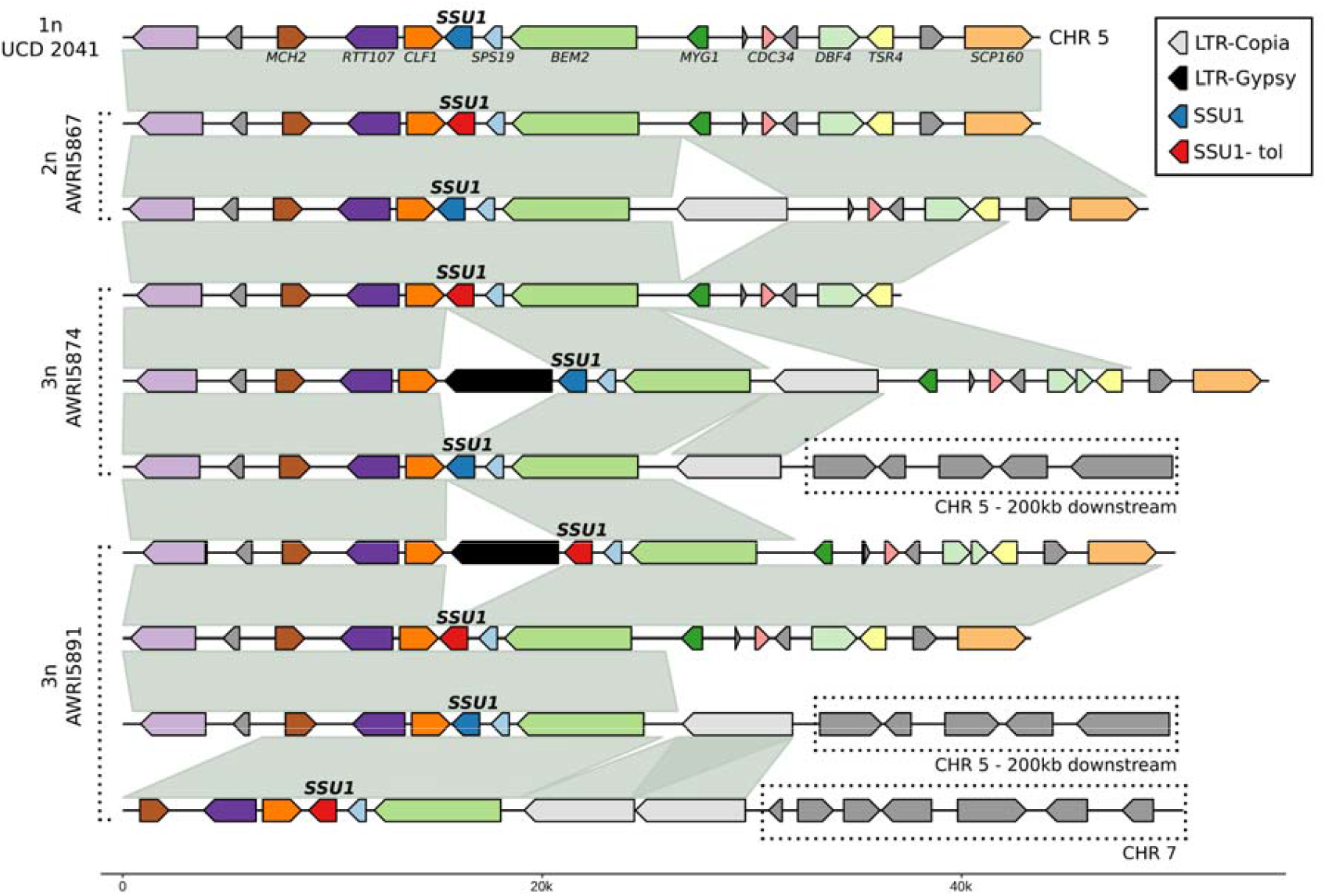
The *SSU1* locus in *B. bruxellensis*. Synteny between the *SSU1* locus of four *B. bruxellensis* strains with differing ploidy levels. SSU1-tol refers to the H1 sulphite tolerant haplotype of *SSU1* shown in Figure 2. Dotted boxes indicate non-syntenic regions, with their genomic locations mapped relative to the reference strain UCD 2041. Homologous genes between haplotypes are shown in the same colour.

Analysis of non-syntenic regions relative to the UCD 2041 reference genome showed that in the triploid strains AWRI5874 and AWRI5891, an additional *SSU1* allele was located ∼200 kb downstream on chromosome 5. Moreover, in the highly SO_2_-tolerant strain AWRI5891, which contains four copies of *SSU1* (Figure 2), one allele was associated with a chromosomal translocation involving chromosome 7 (Figure 3). The loss of synteny adjacent to retrotransposon insertion sites strongly suggests that these elements contribute to structural variation within this locus.

TEs of the Copia and Gypsy LTR retrotransposon families are semi-autonomous replicative sequences that serve as major generators of SVs in fungi [27-29]. Through their replicative nature, they not only create interspersed duplications of themselves but also trigger rearrangements through ectopic recombination [29, 30]. Comparison of AWRI5874 and AWRI5891, both belonging to the AWRI1499 lineage, revealed that the additional *SSU1* copy in AWRI5891 is associated with the chromosome 5-7 translocation. This finding suggests that recombination or excision activity of the LTR-Copia retrotransposon is linked to the acquisition of the extra *SSU1* copy. Inspection of other triploid SO_2_-tolerant isolates indicated that all carry the chromosome 5-7 *SSU1* translocated locus, with some strains containing only a single LTR-Copia insertion, in contrast to the duplicated insertion observed in AWRI5891 (Figure 3).

To assess the prevalence of the specific retrotransposons identified in the *SSU1* locus across the genome, long-read data from winery isolates were analysed to identify SVs relative to the UCD 2041 reference genome. Insertions and deletions were then queried against these specific transposable elements (TEs) and combined using a pangenome graph approach [22].

Non-reference insertions and deletions of the retrotransposons were detected throughout the genomes of the investigated strains, with triploid strains containing a greater number of variants overall (Figure 4). Insertions were more frequent than deletions, with the LTR-Copia retrotransposon showing higher occurrence across nearly all chromosomes in the triploid strains. A total of 293 (238 Copia, 53 Gypsy and 2 Copia-Gypsy), 310 (250 Copia, 58 Gypsy and 2 Copia-Gypsy) and 45 non-reference (26 Copia, 18 Gypsy, 1 Copia-Gypsy) LTR insertions were identified in strains AWRI5891 (3n), AWRI5874 (3n) and AWRI5867 (2n), respectively. Chromosome 9 contained the largest number of insertions and deletions in all three strains (Figure 4).

**Figure 4.**
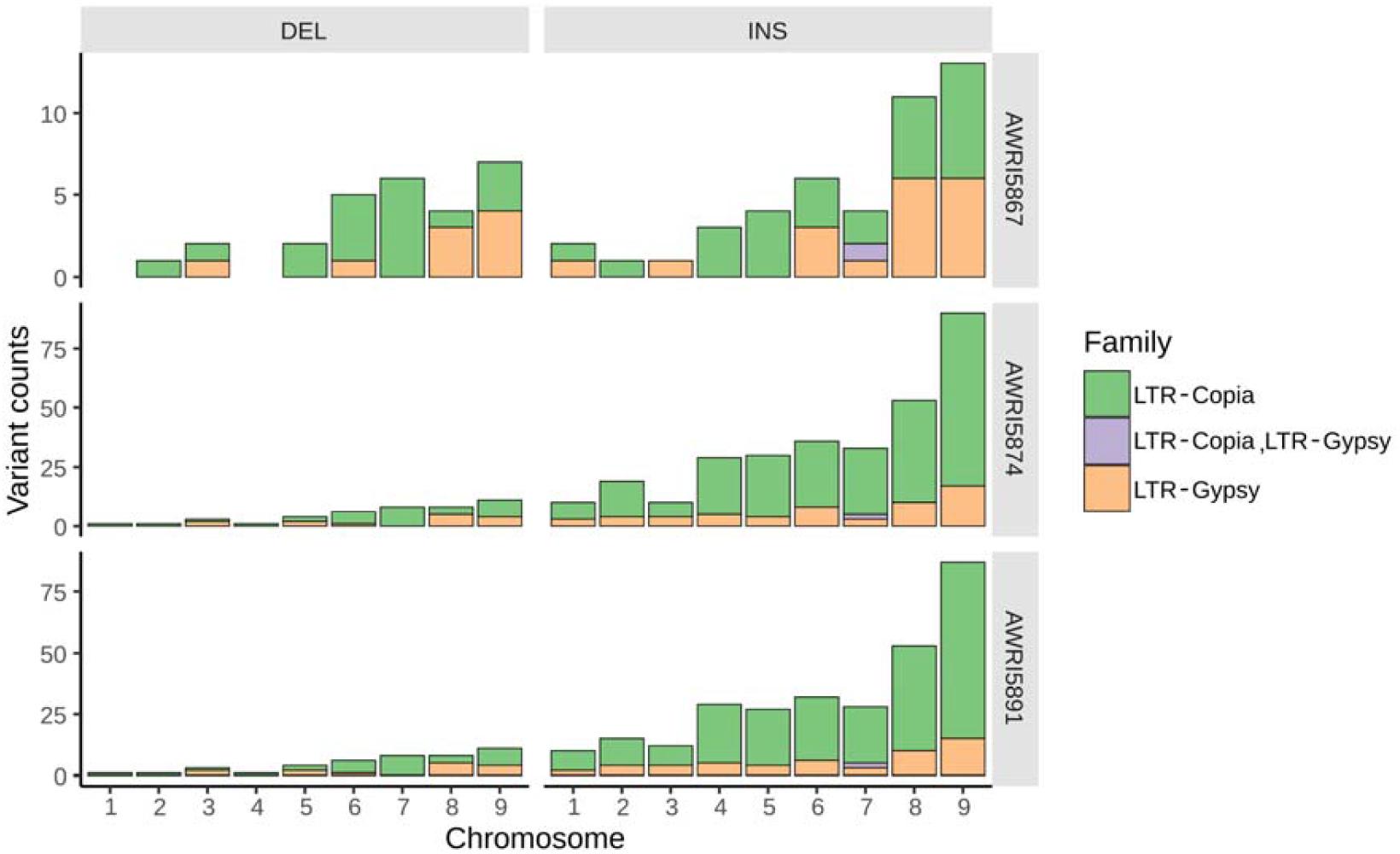
Genome-wide insertions and deletions of the LTR retrotransposons identified within the *SSU1* locus of *Brettanomyces bruxellensis* strains. Bar plots show the total number of variants in strains AWRI5867 (2n), AWRI5874 (3n), and AWRI5891 (3n), identified across each chromosome using the UCD 2041 genome as reference. Only non-reference variations are included. Counts include both complete and partial insertions or deletions of the retrotransposons. LTR-Copia, LTR-Gypsy category refers to specific locations were both types of retrotransposons were identified.

The widespread distribution of these retrotransposons across the genome suggests that they are important drivers of genetic variation in *B. bruxellensis*. At the *SSU1* locus, the presence of these elements indicates that TE-assisted acquisition of *SSU1* copies is the most likely mechanism underlying additional CNVs of this gene. Notably, triploid strains contained a considerably larger number of TE-associated variants, suggesting that polyploid isolates may be more adaptable to environmental pressures [7, 29]. Consistent with this, Bartel, Roach (9) reported more rapid adaptation to SO_2_, accompanied by a higher *SSU1* CNVs, in the triploid strain AWRI1499 compared to the diploid strain AWRI1613.

### 3.3 Adaptive evolution acquired *SSU1* alleles are associated with an LTR-Gypsy retrotransposon

The discovery of retrotransposon insertions within the *SSU1* locus of industry *B. bruxellensis* isolates, together with potential structural variations linked to these mobile elements, suggest a mechanism for the acquisition of additional copies of *SSU1*. Bartel, Roach (9) reported that the triploid strain AWRI1499 acquired extra copies of a specific *SSU1* haplotype following continuous SO_2_ exposure, however, the underlying mechanisms of genomic variation were not investigated. To examine whether retrotransposons might be associated with the reported changes in *SSU1* CNV, previously published nanopore long-read data [9] from AWRI1499 strains subjected to 50 and 100 generations of adaptive evolution in YNB medium, with or without increasing SO_2_ concentrations, were mapped to the *B. bruxellensis* UCD 2041 reference genome (Figure 5). As these data were generated with first-generation nanopore chemistry and had shorter read lengths than the data obtained in the present study, local phasing and assembly of the *SSU1* locus was not possible.

**Figure 5.**
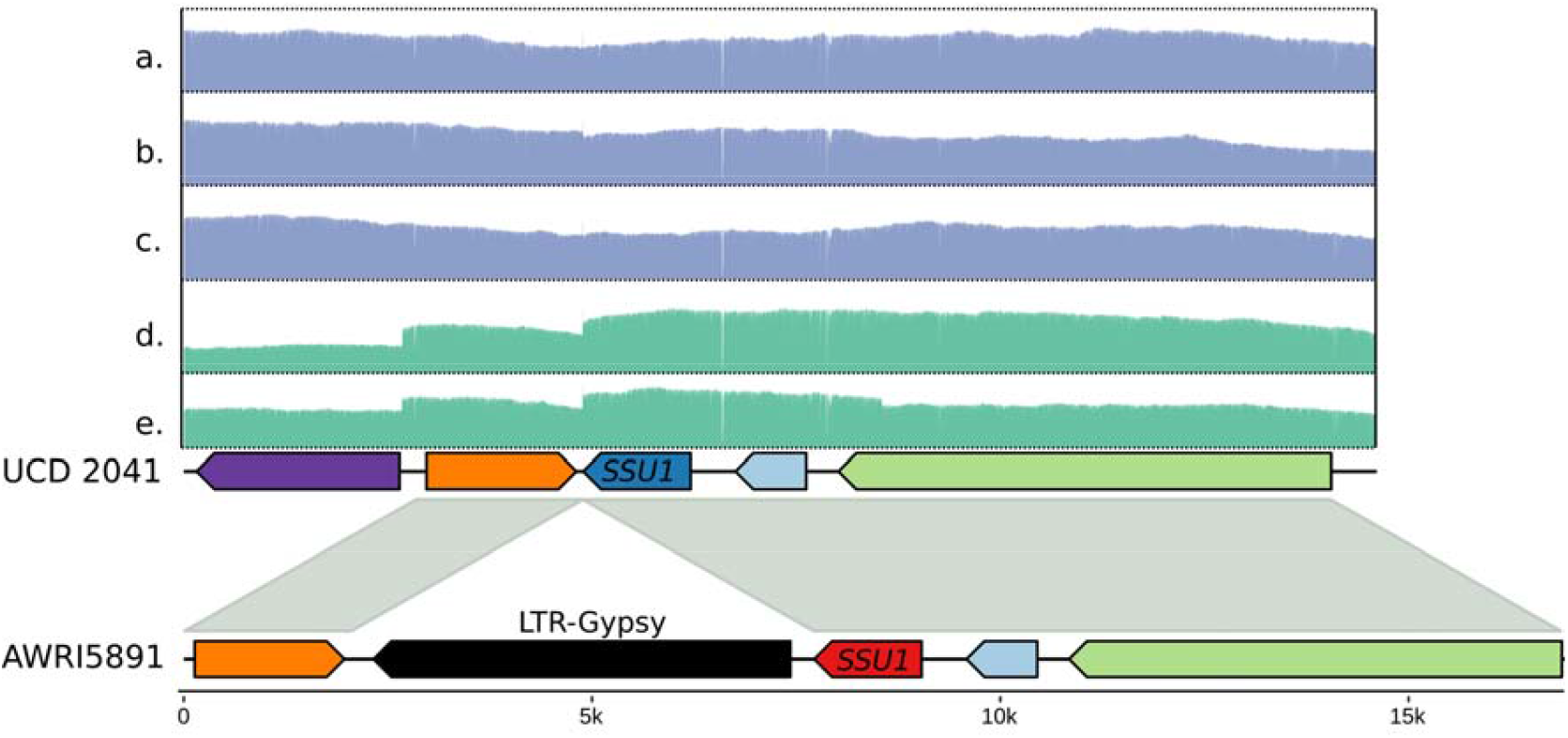
Read depth at the *SSU1* locus in laboratory-evolved *B. bruxellensis* AWRI1499. Previously published long-read datasets for AWRI1499 were mapped to the *B. bruxellensis* UCD 2041 reference genome. Panels show (**a**) parental control (SRR12526261), (**b**) 50 generations in YNB (SRR12526259), (**c**) 100 generations in YNB (SRR12526258), (**d**) 50 generations under increasing sulphite concentrations (SRR12526272), and (**e**) 100 generations under increasing sulphite concentrations (SRR12526271). An increased copy number of the *SSU1* H1 haplotype (Figure 1) was observed in (d and e). A cluster of soft-clipped alignments at *SSU1* coincides with the read-depth shift with all clipped reads containing the Gypsy LTR retrotransposon insertion.

Despite this limitation, read-depth analysis of the *SSU1* locus in these isolates, including the AWRI1499 control, revealed clear shifts in read depth in strains exposed to SO_2_ (Figure 5 d,e). Inspection of the specific site of the read-depth shift identified a structural variant directly downstream of *SSU1*. Synteny analysis with the triploid strain AWRI5891 indicated that this shift coincided with the insertion point of the LTR-Gypsy retrotransposon (Figure 5). Furthermore, examination of soft-clipped regions in reads associated with the increased read depth at *SSU1* revealed that the clipped segments matched the DNA sequence of the LTR-Gypsy retrotransposon identified in industry isolates. Taken together, these results indicate that the additional copies of *SSU1* acquired during adaptive evolution experiments are associated with the LTR-Gypsy retrotransposon allele of *SSU1*.

The metabolic stress imposed by continuous SO_2_ exposure may activate this transposable element and facilitate the acquisition of additional *SSU1* copies, as has been observed in other species [29]. Future studies employing advanced single-cell transcriptomics and long-read sequencing should focus on assessing TE activity under relevant winemaking stress conditions. Identifying the specific stresses that trigger activation of these TEs will be crucial for developing strategies to reduce the emergence of novel *B. bruxellensis* strains with increased SO_2_ tolerance.

## 4. Conclusions

This study demonstrates that variation in SO_2_ tolerance among *B. bruxellensis* isolates from Australian wineries is strongly influenced by ploidy, phylogenetic clade, and structural variation at the *SSU1* locus. High tolerance was consistently associated with duplication of the *SSU1* H1 haplotype, which likely manifest through retrotransposon-mediated structural rearrangements. These results highlight the role of transposable elements as drivers of adaptive genome plasticity in *B. bruxellensi*s, enabling rapid responses by this spoilage yeast to selective pressures imposed by winery practices such as the use of SO_2_. By linking specific haplotype duplications with industry-relevant phenotypes, this work provides a framework for monitoring the emergence of highly tolerant strains.

## 5. Data availability

The sequencing data included in this study are publicly available in the NCBI repository under BioProject PRJNA1330415.

## 6. Conflicts of interest

None declared

## 7. Funding statement

This work was supported by Wine Australia, with levies from Australia’s grape growers and winemakers and matching funds from the Australian Government.

## 8. Acknowledgements

The AWRI is a member of the Wine Innovation Cluster (WIC) in Adelaide. *Brettanomyces bruxellensi*s isolates were provided by Australian wineries.

## Notes

### Competing Interest Statement

The authors have declared no competing interest.

