## Supplementary material for "Retrotransposon-mediated duplication of *SSU1* in high SO_2_ tolerant *Brettanomyces bruxellensis* winery isolates": Figure S1, Table S1

### Supplementary Materials

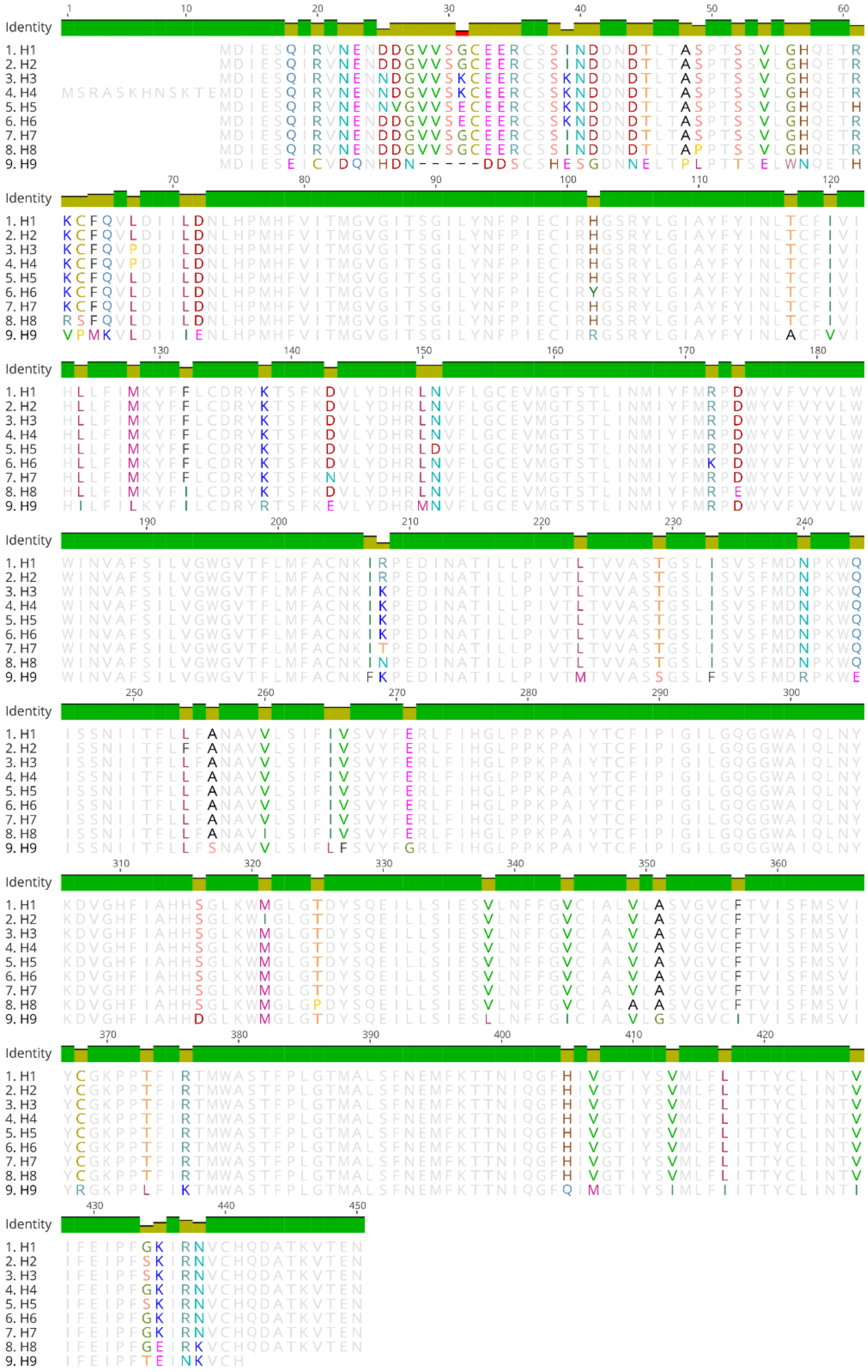


Figure S1. Protein alignment of the nine phased *SSU1* haplotypes identified in *B. bruxellensis* isolates. All mismatches are indicated in the top identity track. DNA sequences encoding the first 12 residues are conserved across all haplotypes but were not annotated in the sequenced isolates. For consistency with Varela, Bartel (14) the first 12 residues of haplotype H4 are included.

Table S1. Description of *B. bruxellensis* isolates included in this study.

| strain | source | ploidy | SO2 tolerance | LTR TE in *SSU1* locus | phylogenetic clade |
|---|---|---|---|---|---|
| AWRI5855 | wine | 3n | high | yes | AWRI1499 |
| AWRI5857 | wine | 3n | high | yes | AWRI1499 |
| AWRI5859 | wine | 3n | high | yes | AWRI1499 |
| AWRI5863 | wine | 3n | high | yes | AWRI1499 |
| AWRI5864 | wine | 3n | high | yes | AWRI1499 |
| AWRI5866 | wine | 3n | high | yes | AWRI1499 |
| AWRI5867 | wine | 2n | low | yes | Wine 3 |
| AWRI5871 | wine | 3n | medium | yes | AWRI1499 |
| AWRI5872 | wine | 2n | low | yes | Wine 3 |
| AWRI5874 | wine | 3n | medium | yes | AWRI1499 |
| AWRI5875 | wine | 3n | medium | yes | AWRI1499 |
| AWRI5876 | wine | 2n | low | no | Wine 3 |
| AWRI5878 | wine | 2n | low | yes | Wine 3 |
| AWRI5880 | wine | 2n | low | no | Wine 3 |
| AWRI5881 | wine | 3n | medium | yes | AWRI1499 |
| AWRI5883 | wine | 2n | low | no | Wine 3 |
| AWRI5888 | wine | 3n | medium | no | Beer |
| AWRI5890 | wine | 3n | high | yes | AWRI1499 |
| AWRI5891 | wine | 3n | high | yes | AWRI1499 |
| AWRI5895 | wine | 3n | medium | yes | AWRI1499 |
| AWRI5896 | wine | 3n | high | yes | AWRI1499 |
| AWRI5898 | wine | 3n | high | yes | AWRI1499 |
| AWRI5900 | wine | 3n | high | yes | AWRI1499 |
| AWRI5901 | wine | 3n | high | yes | AWRI1499 |
| AWRI5903 | wine | 3n | medium | no | Beer |
| AWRI5909 | wine | 2n | medium | no | Kombucha |
